# Clonal evaluation of prostate cancer molecular heterogeneity in biopsy samples by dual immunohistochemistry and dual RNA *in situ* hybridization

**DOI:** 10.1101/818880

**Authors:** Pavithra Dedigama-Arachchige, Shannon Carskadon, Jia Li, Ian Loveless, Mohamed Alhamar, James O. Peabody, Hans Stricker, Dhananjay A. Chitale, Craig G. Rogers, Mani Menon, Tarek A. Bismar, Nilesh S. Gupta, Sean R. Williamson, Nallasivam Palanisamy

## Abstract

Prostate cancer is frequently multifocal. Although there may be morphological variation, the genetic underpinnings of each tumor are not clearly understood. To assess the inter and intra tumor molecular heterogeneity in prostate biopsy samples, we developed a combined immunohistochemistry and RNA in situ hybridization method for the simultaneous evaluation of *ERG, SPINK1, ETV1*, and *ETV4*. Screening of 601 biopsy cores from 120 consecutive patients revealed multiple alterations in a mutually exclusive manner in 37% of patients, suggesting multifocal tumors with considerable genetic differences. Furthermore, the incidence of molecular heterogeneity was higher in African Americans patients compared to Caucasian American patients. About 47% of the biopsy cores with discontinuous tumor foci showed clonal differences with distinct molecular aberrations. ERG positivity occurred predominantly in low Gleason grade cancer, whereas *ETV4* expression was observed mostly in high Gleason grade cancer. Further studies revealed correlation between the incidence of molecular markers and clinical and pathologic findings, suggesting potential implications for diagnostic pathology practice, such as defining dominant tumor nodules and discriminating juxtaposed but molecularly different tumors of different grade patterns.

## INTRODUCTION

Prostate cancer is a heterogeneous disease with varying molecular aberrations observed among patient sub-groups.(1, 2) Of the many molecular alterations present in prostate cancer, E26 transformation-specific (ETS) family gene rearrangements are the most common, occurring in 50-60% of patients.(3) Additionally, another 5-10% of patients are reported to have *SPINK1* over-expression (4), whereas 1-2% display *RAF* kinase gene fusions.(5) Recently, we reported the identification of a pseudogene associated recurrent gene fusion *KLK4-KLKP1* present predominantly in ERG fusion positive tumors (6). Emerging evidence suggests that distinct molecular aberrations have distinct functional roles in prostate cancer and potentially implicated in varying clinical outcomes.(4, 7, 8) ETS gene fusions along with *PTEN* loss has been linked with aggressive prostate cancer.(9) Additionally, *SPINK1* over-expression has been reported to be associated with advanced disease.(10) Although *ERG* and *ETV1* belong to the ETS family of genes, they are known to have distinct functional roles in prostate cancer development.(7) Currently, prostate cancer management options including active surveillance and treatment decisions are governed by pathological observations such as Grade Group (Gleason grade) and tumor volume observed in the initial biopsy samples. However, the association between molecular aberrations and prostate cancer outcome suggests that there may be an untapped role for integration of molecular markers in diagnosis and prognosis. Accordingly, the ETS gene fusion, *TMPRSS2-ERG* and the non-coding RNA, *PCA3* have been explored as diagnostic markers for prostate cancer in efforts to reduce the false positives encountered with PSA.(11, 12) In addition to assisting prostate cancer management decisions, molecular analysis may enable novel therapeutic approaches. In preclinical studies, the use of *SPINK1* screening has been considered for potential therapies using anti-SPINK1 monoclonal antibodies(13) and/or anti-EGFR antibodies.(14) For example, MEK inhibitors have been proposed for patients positive for RAF kinase gene fusions.(5) Moreover, studies targeting ERG gene fusions have been reported.(15-17) Therefore, enabling the evaluation of molecular heterogeneity at the initial biopsy level is an unmet clinical step towards understanding the impact of screening molecular markers in biopsy samples to elucidate tumor heterogeneity in clinical decision making for active surveillance eligibility or other treatment options.

A considerable percentage of prostate cancer patients present with multifocal disease.(18, 19) In current clinical practice, generally the dominant tumor nodule with the largest volume, usually also corresponding to the highest grade and stage tumor, is assumed to drive disease progression.(20) However, additional secondary tumor foci could be clonally different at molecular level and may carry independent driver mutations, which may be associated with cancer progression and the development of metastatic disease (21-23). For example, we have encountered occasional tumors with an admixed high-grade and low-grade component, raising the question of whether this represents “collision” of two clonally different tumors (which should be assigned separate grades), or a single tumor with heterogeneous patterns. Therefore, when making treatment decisions, assessment of independent tumor foci containing identical or different molecular phenotypes may have significant clinical impact, such as for definition of dominant tumors and grading. Determining tumor volume or size in biopsy specimens with discontinuous remains a subject of debate (24-26). In general, a common approach is to assume that discontinuous foci in a single biopsy represent a large, irregularly-shaped tumor.(27, 28) However, some data support clonally different tumors.(26) Given that tumor volume percentage is a critical parameter used in the selection of prostate cancer management options such as active surveillance, enabling the assessment of clonal differences in the foci in cores with discontinuous foci may be important. A recent study reported that the cancer incidence and prognosis vary according to the location of the tumors within in the prostate.(29) However, another study found no advantage of zonal location of cancer over other prognostic factors.(30) These studies were based on morphological evaluation only. Therefore, determining whether tumor location in the prostate has any specific association with particular genetic aberrations is required. Keeping the above points in view and considering the importance of differentiating inter and intra tumor heterogeneity, we carried out a comprehensive analysis, evaluating the incidence of several recurrent prostate cancer molecular markers in prostate needle biopsy samples to facilitate a better understanding of molecular heterogeneity and the clonal nature of multifocal disease with a focus on racial differences.

## Materials and methods

### Study design and patient selection

A total of 601 biopsy cores were collected from 120 patients who had undergone ultrasound-guided transrectal needle biopsy procedures from July 2016 to October 2016 in the Henry Ford Health System (Detroit). The pathological reports of the needle biopsies were reviewed and the location, Gleason grade (including Grade Group), and the tumor volume percentage of the prostate cores were recorded. The presence of discontinuous foci was determined as previously described.(26) Biopsy cores containing benign tissue, high grade intraepithelial neoplasia, atypical glands / atypical small acinar proliferation, and varying Gleason patterns for each patient were collected for further evaluation. The patient age, race, family history of cancer, initial PSA value (PSA value closest to the study biopsy), status of additional needle biopsies, subsequent radical prostatectomy, subsequent radiation and/or hormone treatment, and the last PSA (most recent PSA value recorded after study biopsy) were documented. In all cases, informed consent and Institutional Review Board approval were obtained.

### Dual RNA in situ Hybridization and Dual Immunohistochemistry

Given the limited availability of biopsy tissues, we developed a novel four-color multiplex assay for the simultaneous evaluation of *ERG, SPINK1, ETV1* and *ETV4*. Due to the lack of cancer specific antibodies for *ETV1* and *ETV4*, dual RNA in situ hybridization was performed first for *ETV1* and *ETV4* followed by dual immunohistochemistry for ERG and SPINK1 on the same tissue section. Specifically, slides were incubated at 60° C for 1 hour. Tissues were then deparaffinized by immersing in xylene twice for 5 minutes each with periodic agitation. The slides were then immersed in 100% ethanol twice for 3 minutes each with periodic agitation, then air-dried for 5 minutes. Tissues were circled using a pap pen (Vector, H-4000), allowed to dry, and treated with H2O2 for 10 minutes. Slides were rinsed twice in distilled water, and then boiled in 1X Target Retrieval for 15 minutes. Slides were rinsed twice in distilled water, and then treated with Protease Plus for 15 minutes at 40° C in a HybEZ Oven (Advanced Cell Diagnostics, 310010). H2O2, 1X Target Retrieval, and Protease Plus are included in the RNAscope Pretreatment kit (Advanced Cell Diagnostics, 310020). Slides were rinsed twice in distilled water, and then treated with both *ETV1* (Advanced Cell Diagnostics, 311411), and *ETV4* (Advanced Cell Diagnostics, 478571-C2) probes at a 50:1 ratio for 2 hours at 40°C in the HybEZ Oven. Slides were then washed in 1X Wash Buffer (Advanced Cell Diagnostics, 310091) twice for 2 minutes each. Slides were then stored overnight in a 5X SSC solution. The next day, slides were again washed in 1X Wash Buffer twice for 2 minutes each. Slides were then treated with Amp 1 for 30 minutes, Amp 2 for 15 minutes, Amp 3 for 30 minutes, and Amp 4 for 15 minutes, all at 40° C in the HybEZ oven with 2 washes in 1X Wash Buffer for 2 minutes each after each step. Slides were then treated with Amp 5 for 30 minutes and Amp 6 for 15 minutes at room temperature in a humidity chamber with 2 washes in 1X Wash Buffer for 2 minutes each after each step. Red color was developed by adding a 1:60 solution of Fast Red B: Fast Red A to each slide and incubating for 10 minutes. Slides were washed in 1X Wash Buffer twice for 2 minutes each, then treated with Amp 7 for 15 minutes and Amp 8 for 30 minutes at 40° C in the HybEZ oven with 2 washes in 1X Wash Buffer for 2 minutes each after each step. Slides were then treated with Amp 9 for 30 minutes and Amp 10 for 15 minutes at room temperature in a humidity chamber with 2 washes in 1X Wash Buffer for 2 minutes each after each step. Brown color was developed by adding a solution of Betazoid DAB (1 drop DAB to 1ml Buffer; Biocare Medical, BDB2004L) to each slide and incubating for 10 minutes. Amps 1-10 and Fast Red are included in the RNAscope 2.5 HD Duplex Detection Reagents (Advanced Cell Diagnostics, 322500). Slides were washed twice in distilled water, and then washed in 1X EnVision FLEX Wash Buffer (DAKO, K8007) for 5 minutes. Slides were then treated with Peroxidazed 1 (Biocare Medical, PX968M) for 5 minutes and Background Punisher (Biocare Medical, BP974L) for 10 minutes with a wash of 1X EnVision FLEX Wash Buffer for 5 minutes after each step. Anti-ERG (EPR3864) rabbit monoclonal primary antibody (1:50; Abcam, ab92513) and a mouse monoclonal against SPINK1 (1:100; Novus Biologicals, H00006690-M01) were added to each slide, which were then cover slipped with parafilm, placed in a humidifying chamber, and incubated overnight at 4° C. The next day, slides were washed in 1X EnVision Wash Buffer for 5 minutes and then incubated in Mach2 Doublestain 1 (Biocare Medical, MRCT523L) for 30 minutes at room temperature in a humidifying chamber. Slides were then rinsed in 1X EnVision Wash Buffer 3 times for 5 minutes each. Slides were then treated with a Ferangi Blue solution (1 drop to 2.5ml buffer; Biocare Medical, FB813S) for 7 minutes, washed in 1X EnVision FLEX Wash Buffer for 5 minutes, and then treated with a Vina Green solution (1 drop to 1ml buffer; Biocare Medical, BRR807AS) for 15 minutes. Slides were then rinsed 2 times in distilled water, then treated with EnVision FLEX Hematoxylin (DAKO, K8008) for 2 minutes. Slides were rinsed several times in distilled water, immersed in a 0.01% ammonium hydroxide solution, and then rinsed twice in distilled water. Slides were then dried completely. Slides were dipped in xylene approximately 15 times. EcoMount (Biocare Medical, EM897L) was added to each slide, which was then cover slipped. The staining of the whole-mount radical prostatectomy case was carried out using a modified procedure described previously (31).

### Statistical analysis

Two-sample t-test was used to study the association between molecular marker expression and patient age, initial PSA, last PSA and the duration of time between the study biopsy and subsequent treatment. Cox proportional hazards model was used to analyze the association of molecular marker expression with the subsequent treatment, radical prostatectomy and radiation. The cases where subsequent treatment information was not available were treated as censored. In all other cases, Pearson’s chi-square test was used. In all analyses, P-values < 0.05 were considered statistically significant.

## RESULTS

We collected 601 biopsy cores from 120 consecutive patients (Caucasian American, 67; African American, 47 and 6 from other racial groups, (**Table S1**) who had undergone prostate needle biopsy procedures from July to October 2016. Of the 120 patients, 75 included standard 12 core needle biopsies where tissue was extracted from 12 specific prostate locations, namely, right lateral base, right lateral mid, right lateral apex, right base, right mid, right apex, left lateral base, left lateral mid, left lateral apex, left base, left mid, and left apex (**Figure 1A**). In 39 patients, additional targeted prostate tissue cores had been obtained during the biopsy which were also included in the evaluation for molecular markers. Six patients had 6 or fewer number of prostate locations sampled during the needle biopsy. Due to the limited availability of biopsy tissue in some of the tissue blocks, we were not able to evaluate all biopsy cores with cancer in some patients. The number of biopsy cores collected from each patient ranged from 1 to 13 with a median of 4 cores. Overall, 572 biopsy cores originated from standard 12 core biopsy locations, whereas 29 cores had been collected from other prostate locations (**Figure 1A, Table S2**). Biopsy cores selected for evaluation included, high grade intraepithelial neoplasia, atypical/atypical small acinar proliferation and Gleason graded prostate cancer samples (Grade Groups 1-5, **Figure 1B**). Out of the 601, 119 cores were found to have discontinuous foci, where tumor foci occurred in the same biopsy core separated by benign tissues.

**Figure 1:**
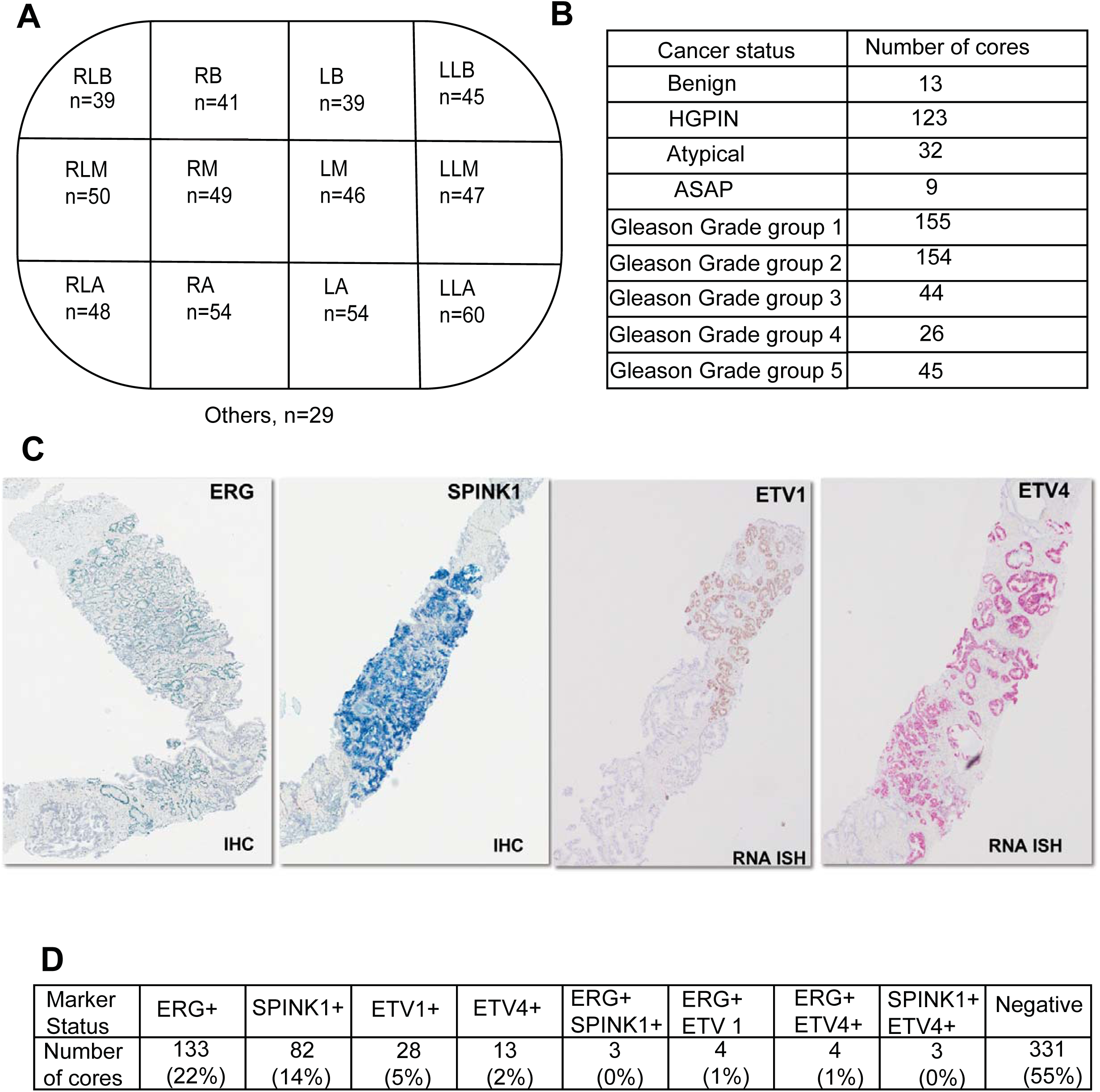
The expression of multiple prostate markers in prostate needle biopsies. (A) The cancer status of the needle biopsy cores used in the study. HGPIN= high grade intraepithelial neoplasia, Atypical= atypical/atypical small acinar proliferation) (B) The locations of the prostate from where the needle biopsy cores were obtained. The number of cores originating from each location is shown. RLB =right lateral base, RLM = right lateral mid, RLA = right lateral apex, RB = right base, RM = right mid, RA = right apex, LLB = left lateral base, LLM = left lateral mid, LLA = left lateral apex, LB = left base, LM = left mid, LA = left apex. (C) The expression of ERG, SPINK1, *ETV1* and *ETV4* evaluated by dual immunohistochemistry and dual RNA in situ hybridization. The immunohistochemistry and RNA in situ hybridization signals observed for ERG (green), SPINK1 (blue), *ETV1* (brown) and *ETV4* (red) in representative needle biopsy cores are shown. (D) The expression profile of ERG, SPINK1, *ETV1* and *ETV4* in the needle biopsy core cohort. The percentage of cores with each expression profile is also shown.

We performed dual RNA in situ hybridization for *ETV1* and *ETV4* and subsequent dual immunohistochemistry for ERG and SPINK1, on the same tissue section, on the 601 biopsy cores collected (**Figure 1C**). Of the 601 cores evaluated, 270 cores (45%) were found to be positive for at least one of the molecular markers tested (**Figure 1D**). Among the positive cores, ERG was found to be the most prominent marker with positivity observed in a total of 144 cores (24%), followed by SPINK1 positivity in 88 (15%). *ETV1* and *ETV4* were positive in 32 (5%) and 20 (3%) cores, respectively. Importantly, 14 (2%) cores showed the expression of two different molecular markers (ERG+/SPINK1+; ERG+/*ETV1*+; ERG+/ *ETV4*+ and SPINK1+/ *ETV4*+) in the same biopsy core but in two different tumor foci. Notably, all cores with dual marker expression occurred on cores with discontinuous tumor foci where the expression of the two molecular markers was mutually exclusive and was noted in separate tumor foci, indicating different clonal origin of the tumor foci. We did not observe any cores with positivity for more than one molecular marker in a single focus. Of note, a significant number of tumor foci (n=331) were negative for all the four molecular markers evaluated.

Next, we analyzed the expression profile of ERG, SPINK1, *ETV1* and *ETV4* in needle biopsy cores carrying discontinuous tumor foci. Overall, 57 (48%) cores with discontinuous tumor foci showed the expression of a single molecular marker in all tumor foci (**Figure 2A, Figure 2B**), suggesting similar clonal origin of the discontinuous foci. Additionally, 6 cores with discontinuous foci showed negative results for all the tested molecular markers in both tumor foci (**Figure 2A, Figure 2C**). Notably, 56 (47%) cores demonstrated discordant molecular marker expression in separate tumor foci within the same core biopsy (**Figure 2A**), suggesting different clonal origin of the discontinuous foci. Among the cores showing discordant molecular marker expression, 14 displayed mutually exclusive expression of two molecular markers in separate tumor foci as described before (**Figure 1D, Figure 2A, Figure 2C**). The rest of the cores showed the expression of a single molecular marker in one focus, whereas the other focus was negative for all four markers (**Figure 2A, Figure 2E**).

**Figure 2:**
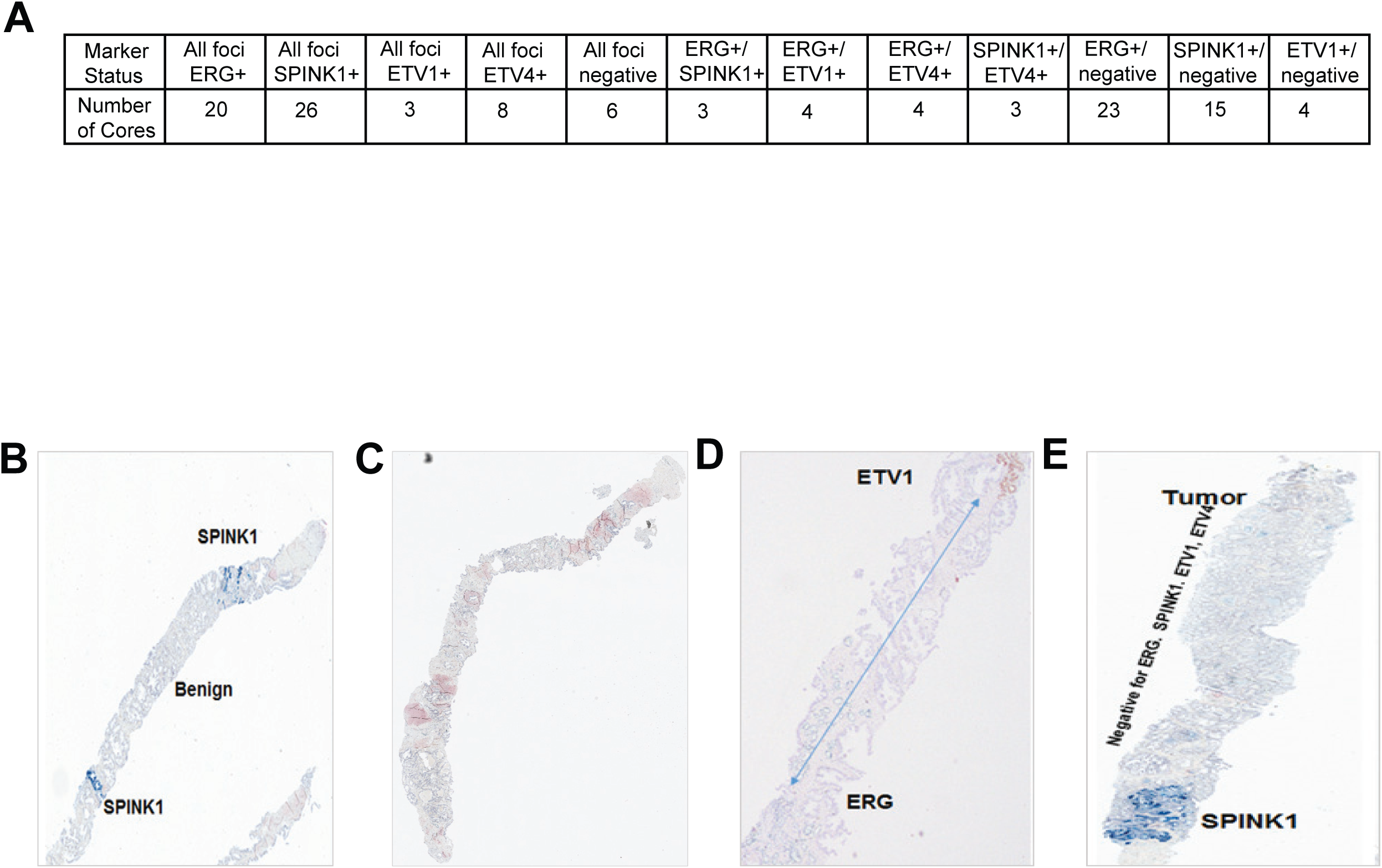
The expression of ERG, SPINK1, *ETV1* and *ETV4* in prostate needle biopsy cores with discontinuous tumor foci. (A) The expression profile of cores with discontinuous tumor foci. For example, all foci ERG+ refers to cores with ERG expression in all the distinct tumor foci. All foci negative refers to cores with no expression of the tested markers in the tumor foci. ERG+/SPINK1+ refers to cores showing mutual exclusive expression of ERG and SPINK1 in distinct tumor foci. ERG+/negative refers to cores where ERG expression was observed only in some tumor foci with other tumor foci being negative for all the tested markers. (B) The expression of SPINK1 in all of the discontinuous tumor foci. The benign tissue separating the two distinct tumor foci is noted. (C) The absence of ERG, SPINK1, *ETV1* and *ETV4* in all of the tumor foci. (D) The mutually exclusive expression of ERG and *ETV1* in discontinuous tumor foci in the same biopsy core. (E) The expression of SPINK1 in one tumor foci while the rest of the tumor foci remains negative for the molecular markers.

We then studied the association of ERG, SPINK1, *ETV1* and *ETV4* expression with the corresponding cancer status of the needle biopsy cores. Of the 601 needle biopsy cores, 424 cores included cancer graded from Grade Group 1 to 5 (**Figure 1B**). Of the 424 cores having Grade Group 1-5 cancer, 242 cores (57%) were positive for at least one marker or more than one (**Table 1**). In contrast, out of the 123 cores with high grade intraepithelial neoplasia, only 12 cores (10%) were positive for any one of the four markers, whereas all 13 benign cores were negative. Statistical analysis confirmed that the expression of molecular markers is preferentially associated with grade group 1-5 cancer compared to both benign (p<0.001) and high grade intraepithelial neoplasia tissues (p<0.001). Additionally, only 16 cores of the 41 cores (39%) with atypical/atypical small acinar proliferation were positive for at least one molecular marker. The expression of molecular markers in atypical/atypical small acinar proliferation cores were significantly higher compared to benign and high grade intraepithelial neoplasia tissues (p<0.001). Further analysis revealed significant expression of ERG in lower Grade Groups (grade groups 1 and 2) compared to high Grade Groups (Grade Groups 3-5, p<0.001). On the contrary, *ETV4* expression was seen more in higher Grade Group samples (Grade Groups 3-5, p=0.04). No significant association was seen between Grade Group and the expression of SPINK1 or *ETV1* or any of the dual markers.

As a further step, we also explored, if there is any relationship between cancer status and the locations of the tumors in the prostate. For this analysis, we used 572 biopsy cores that were extracted from the 12-standard biopsy locations of the prostate (**Figure 1A, Table S2**). The incidence of Grade Group 1-5 cancers, atypical/atypical small acinar proliferation foci in each of the 12 core locations ranged from 67% to 89% (**Figure 3, Table S3**). Interestingly, statistical analysis showed that Grade Group 1-5 tumors, and atypical/atypical small acinar proliferation occurred significantly more in the left side of the prostate (81%) compared to the right side (74%, p=0.03). Furthermore, out of all the 12 locations, left apex showed a significantly higher percentage of Grade Group 1-5 cancer, and atypical/atypical small acinar proliferation (89%, p=0.04). In contrast, the percentages of graded cancer, atypical/atypical small acinar proliferation cases observed with other prostate locations were not significantly different from each other (p>0.05). We also studied, if any of the individual Grade Groups, low Grade Group (Grade Groups 1 and 2) or high Grade Group (Grade Groups 3, 4 and 5) associate with any specific prostate location. However, no statistically significant association was observed.

**Figure 3:**
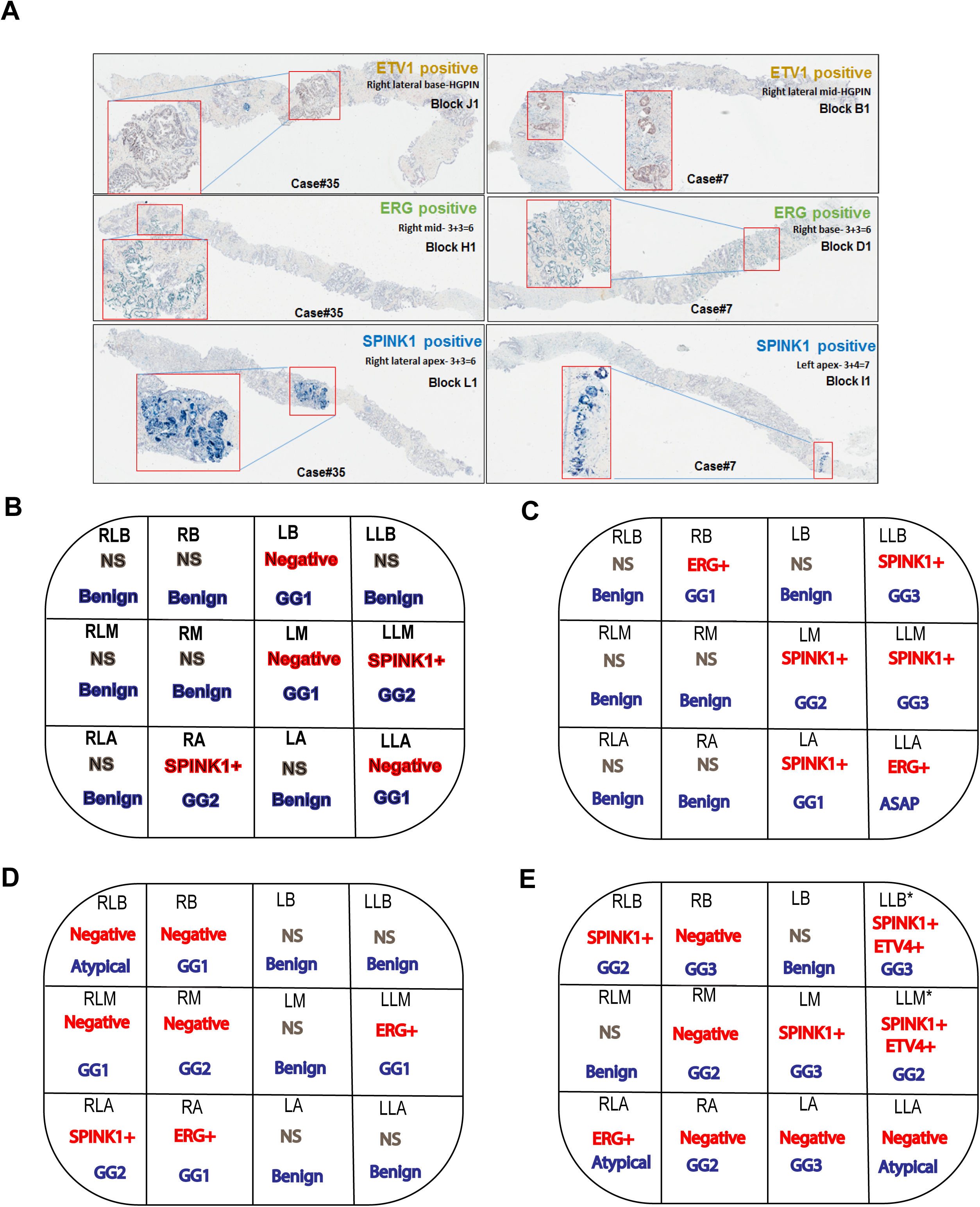
The cancer status of the cores obtained from different locations of the prostate. RLB = right lateral base, RLM = right lateral mid, RLA = right lateral apex, RB = right base, RM = right mid, RA = right apex, LLB = left lateral base, LLM = left lateral mid, LLA = left lateral apex, LB = left base, LM = left mid, LA = left apex.

Next, we studied the association between the incidence of molecular markers and tumor location in the prostate by analyzing the expression profile of ERG, SPINK1, *ETV1* and *ETV4* observed in the 572 biopsy cores originating from the 12 prostate locations (**Figure 4, Table S4**). The percentage of cores showing expression for at least one molecular marker ranged from 33% to 54% in the 12 core locations. Specifically, ERG showed significantly higher expression in the left side of the prostate (29%) compared to the right side (20%, p=0.02) without any association for a specific location on the left side. On the contrary, *ETV1* displayed significantly more expression on the right side (8%) than the left (2%, p=0.00). Furthermore, *ETV1* occurred most in the right lateral mid (10%) compared to the rest of the core locations (4%, p=0.04). SPINK1 and *ETV4* expression did not show any association with a particular location in the prostate.

**Figure 4:**
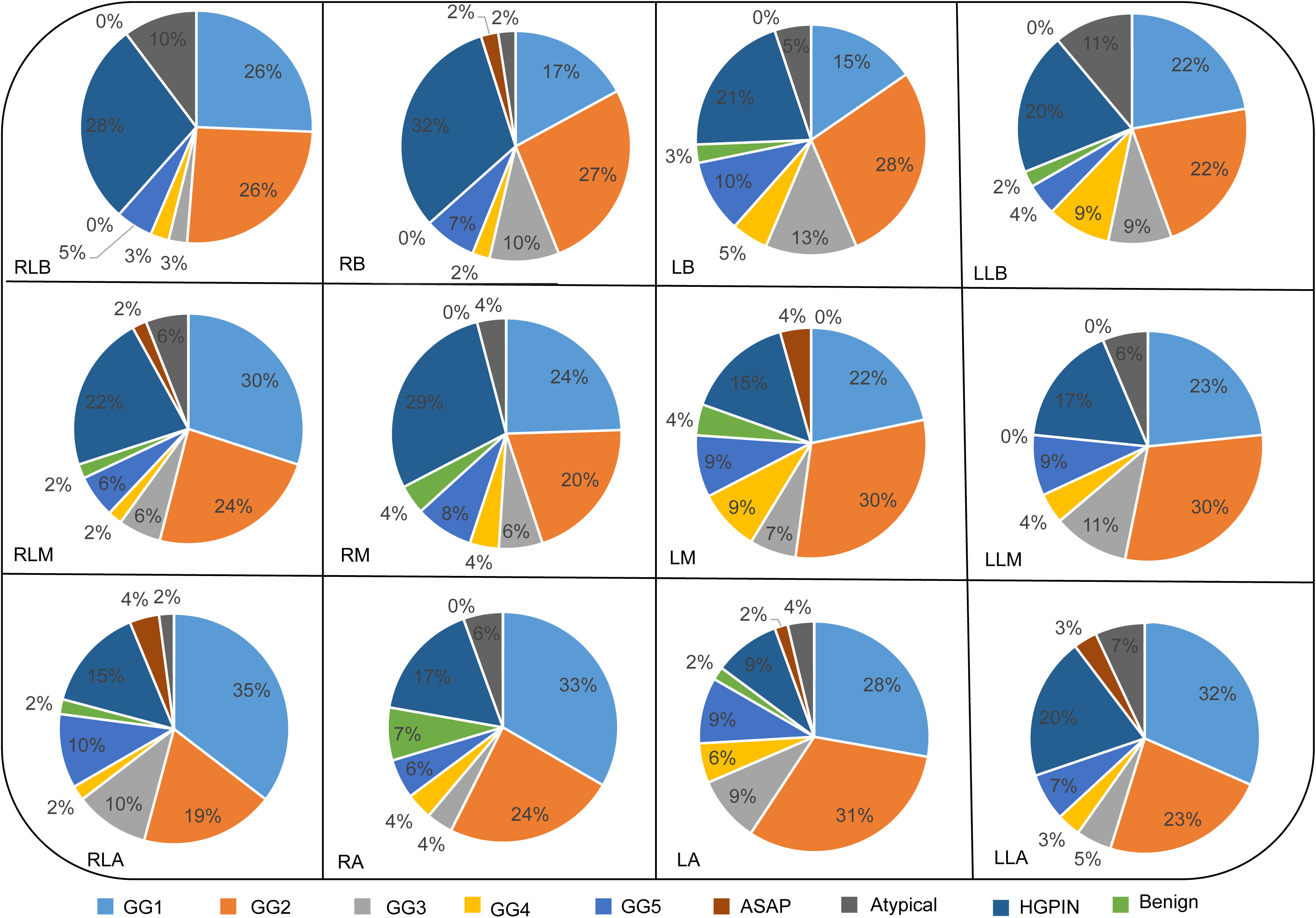
The expression of molecular markers in cores obtained from different locations of the prostate. RLB = right lateral base, RLM = right lateral mid, RLA = right lateral apex, RB = Right base, RM = Right mid, RA = Right apex, LLB = Left lateral base, LLM = Left lateral mid, LLA = Left lateral apex, LB = Left base, LM = Left mid, LA = left apex.

Then we explored the overall incidence of ERG, SPINK1, *ETV1* and *ETV4* in our patient cohort. Altogether, 42 (35%) patients in the cohort tested negative for all 4 molecular makers (**Table S5**). Of the rest, 53 (44%) patients, which included 32 Caucasian American and 18 African Americans, showed the expression of at least one marker. A total of 23 patients (19%), including 10 Caucasian American and 12 African American patients showed positivity for two different markers. Two patients (2%), both African American, were positive for ERG, SPINK1 and *ETV1* in three different cores, indicating the diverse molecular subtypes present in prostate cancer. Overall, 78 (65%) patients showed the expression of at least one molecular marker. Of these, ERG expression was observed in 52, whereas SPINK1 was positive in 29, *ETV1* in 19, and *ETV4* in 8. In agreement with previous studies, statistical analysis showed significantly more positivity for SPINK1 in African Americans (p<0.01) compared to Caucasian Americans. Additionally, *ETV4* expression was also significantly higher in African Americans (p= 0.04), whereas ERG and *ETV1* expression was not significantly different between the two races.

To obtain a comprehensive understanding of molecular heterogeneity and the clonal nature of the disease detected at the biopsy level, we then looked at the expression of ERG, SPINK1, *ETV1* and *ETV4* in the patient cohort in detail. Specifically, 76 (63%) patients showed either the absence of all four molecular markers or the expression of a single molecular marker across all the tested biopsy cores, suggesting either disease arising from a single clonal origin or the presence of hitherto unidentified driver molecular markers in prostate cancer (**Table 2**). The rest of the 44 (37%) cases displayed distinct marker status across biopsy cores, suggesting cancer originating from multiple clones. Of these, some patients showed the occurrence of a single molecular marker in some biopsy cores, whereas the rest of the biopsy cores tested negative for the molecular markers (**Table 2, Figure 5A**). Other patients displayed even more complex expression patterns with multiple molecular markers being seen across different biopsy cores (**Table 2, Figures 5B-5E**), indicating extensive molecular heterogeneity in tumors arising at different locations of the prostate. Interestingly, statistical analysis showed that the incidence of distinct marker status across biopsy cores is significantly higher in African Americans compared to Caucasian Americans (P=0.025). Of note, as previously described (**Table S5**), the two patients in the cohort who displayed the expression of three molecular markers in three different biopsy cores from three different locations of the prostate (**Figure 5E**) were both African American. Overall, our results highlight the existence of marked inter tumor molecular heterogeneity in a subset of cases with multi focal cancer.

**Figure 5:**
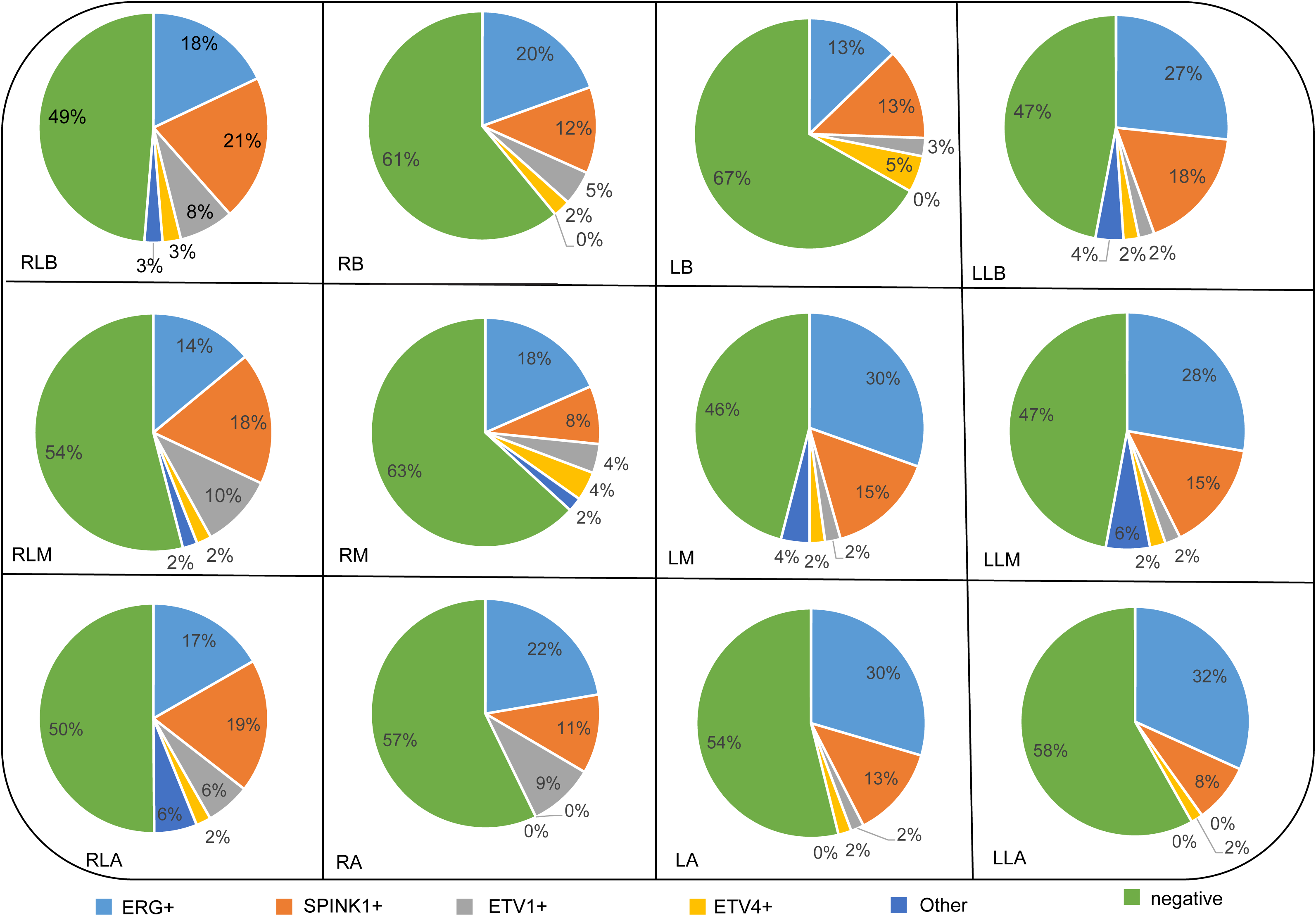
Inter tumor molecular heterogeneity observed in the patient cohort. (A) Representative patient case showing molecular heterogeneity with some tissue cores positive for *SPINK1*, while the other Graded cancer cores are negative for *ERG, SPINK1, ETV1* and *ETV4*. The cancer status of the tissue cores are also noted. NS = tissue cores not screened. GG = Grade Group. Atypical = atypical/atypical small acinar proliferation. (B) Representative patient case showing molecular heterogeneity with *ERG* and *SPINK1*. (C) Representative patient case showing molecular heterogeneity with *ERG* and *SPINK1* while other Graded cancer cores are negative for all molecular markers screened (D) Representative patient case showing molecular heterogeneity with ERG, SPINK1 and *ETV1*. Some Graded tissue scores are negative for all four molecular markers. The tissue cores with discontinuous tumor foci are marked with *. (E) Two patient cases showing the expression of multiple molecular markers in different needle biopsy cores.

Next, we were interested to study whether the biopsy cores from each patient truly represent all the tumor foci in the prostate or if any of the foci are missed during standard biopsy procedure. We selected a representative case with subsequent radical prostatectomy and compared the tumors in the biopsy with matching whole mount radical prostatectomy tissue by screening for ERG, SPINK1, *ETV1* and *ETV4*. Comparative analysis of needle biopsy results and whole mount radical prostatectomy results revealed that multiple secondary small foci positive for SPINK1 were not represented in the needle biopsy (**Figure 6**). Tumors represented in the biopsy with corresponding topographical location were matched in the radical prostatectomy tissue, although the highest grade area, Gleason score 4+3=7 (Grade Group 3), was not represented in this section from mid-prostate due to relatively small size. In an unpublished study, we evaluated 987 radical prostatectomy wholemount tissue for ERG, SPINK1, *ETV1, ETV4, ETV5* and found several secondary foci positive for molecular markers. The significance of the marker positive secondary tumors are not known.

**Figure 6:**
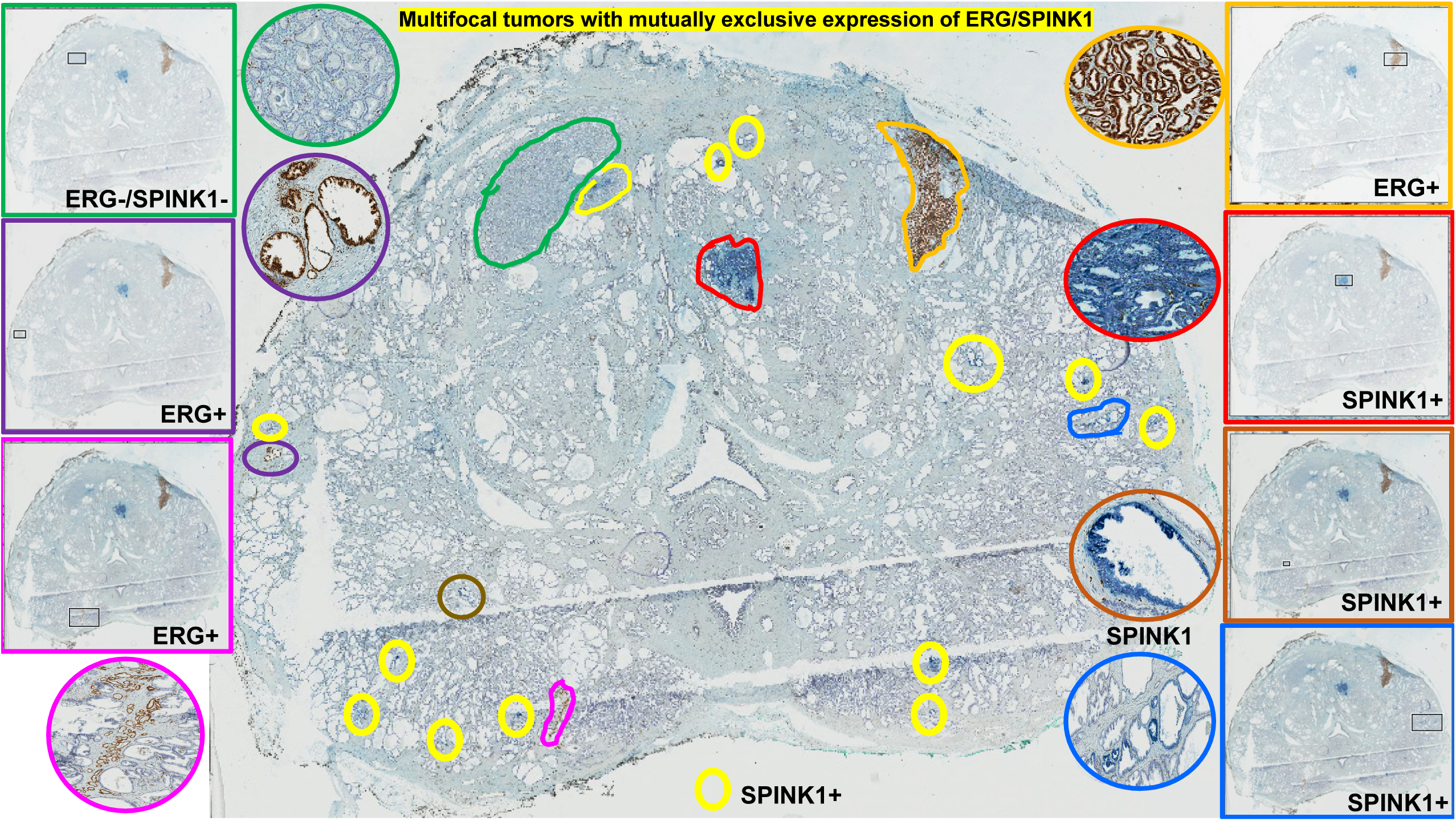
A representative whole-mount radical prostatectomy specimen from one of the cohort patients shows multiple scattered tumor foci with distinct biomarker staining patterns. Largest tumors include those in the left anterior (predominantly negative but with focal adjacent SPINK1 positivity) and right anterior (ERG positive). Scattered isolated foci of SPINK1 positivity are present (yellow circles). Isolated high-grade prostatic intraepithelial neoplasia with ERG positivity is present in the left lateral (purple circles). Other scattered foci are shown topographically in colored boxes, with matching higher magnifications in the corresponding color of circles or ovals.

Finally, we investigated the association of molecular markers with the clinical and pathological parameters. Using t-test, we first studied if any of the molecular markers we screened are associated with the age of patients. The age of the patients in the cohort ranged from 44 to 86 years with an average of 66 years (**Table S1**). Interestingly, we observed that positive expression of SPINK1 (p=0.01) and the presence of *ERG*+*/SPINK1*+ (p=0.02) are associated with young men with prostate cancer (**Table S6**). Mean age observed with SPINK1 positive patients was 62 years while mean age seen for patients with ERG+/SPINK1 was 57 years. The other markers did not show any significant association with patient age. Then we analyzed the association between marker expression and the number of biopsies a patient had undergone. Of the 120 patients, 82 underwent only the study biopsy, 38 (32%) had multiple biopsies, ranging from 2 to 8. Among the patients with multiple biopsies, 25 had at least one biopsy procedure conducted before the 2016 study biopsies, which were used for the evaluation of molecular markers. On the other hand, 17 patients had additional biopsies after the study biopsy (**Table S1**). We used Pearson’s chi-square test to explore the association between molecular marker expression and the number of biopsies. Intriguingly, *ETV1* expression (p=0.01) associated with single prostate biopsies (**Table S7**). No other association was observed between the number of biopsies and the molecular marker expression. We also explored the association of molecular marker expression with initial PSA and the last PSA values of the patients in our cohort using t-test. The initial PSA, which was either collected before the study biopsy or within a month of the study biopsy ranged from 0.026 ng/mL to 1810.8 ng/mL with an average of 23.4 ng/mL. The last PSA values, which were the most recent PSA values recorded after the study biopsy, were available for 112 patients. Of these, 52 patients showed a PSA value of <0.1 ng/mL. In the other 60 patients, the last PSA ranged from 0.2 to 27.2 ng/mL (**Table S1**). No significant associations were observed with either initial PSA (**Table S8**) or last PSA (**Table S9**) for any of the molecular markers. As a further step, we used Cox proportional hazards model to analyze if there is any association between the molecular marker expression and the presence of subsequent treatment. Subsequent to biopsy procedure, twelve patients had received radiation while 4 had been treated with hormone therapy. A total of 51 patients had undergone subsequent radical prostatectomy. Of the 51 cases, 7 patients had hormone or/and radiation therapy after radical prostatectomy. The information on subsequent treatment method was not available for 48 patients (**Table S1**). Interestingly, while the expression of SPINK1 (p= 0.01), *ETV1* (p=<0.01), and ERG+/*ETV1* (p=<0.01) expression associated with the presence of later treatment, the expression of only ERG (p=0.05) in the biopsy specimens associated with the absence of subsequent treatment (**Table S10**). Next, we investigated if any of the molecular markers associate with either subsequent radical prostatectomy or radiation. Here only the patients who underwent radical prostatectomy or radiation as the primary treatment method were considered, and the patients who underwent radiation as salvage therapy were not included. The association of molecular marker expression with hormone therapy was not investigated since the number of patients who received hormone therapy was small. When analyzed by Cox proportional hazards model, we observed that *ETV1* expression (p=<0.01) and ERG*+/ETV1+* expression (p=<0.01) are associated with later radical prostatectomy (**Table S11**). No significant association was observed with the molecular marker expression and subsequent radiation (**Table S12**). Finally, using t-test, we explored if the expression of the markers associated with the duration of time occurred between the study biopsy and the subsequent treatment. No significant association was observed (**Table S13**).

## DISCUSSION

To assess the extent of tumor molecular heterogeneity and multi clonal nature detected at biopsy level, we evaluated the incidence of ERG, SPINK1, *ETV1* and *ETV4* in 601 prostate needle biopsy cores collected from 120 patients. We discovered that a considerable fraction of the cases (37%) display inter-tumor molecular heterogeneity, with different driver alterations present in different biopsy cores from the same patient. Subsets of patients showed the positivity for as much as three molecular markers across different cores, suggesting extensive clonal differences even in single patient cases. Currently, most clinical decisions are made based on the dominant tumor nodule which displays the largest tumor volume, usually also being the highest grade and stage tumor. Occasionally, a smaller volume tumor may be of higher grade or stage, supplanting the largest tumor as the most clinically relevant. However, there are several diagnostic scenarios in which molecular biomarker assessment may play a role in in the future for classifying the dominant tumor, such as with confluent but molecularly different foci of different grade patterns. As an example, it is not unusual to encounter large tumors that would conventionally be graded as Gleason score 3+4=7 (Grade Group 2) or Gleason score 4+3=7 (Grade Group 3), yet in which sizeable areas would appear to be Gleason score 4+4=8 (Grade Group 4). If such foci show different biomarker patterns, it would be logical to grade them separately, in the same way that they would be graded separately if they were anatomically not confluent, especially in view of the exceedingly high rate of multifocality of prostate cancer.

Despite the critical role of tumor volume percentage in determining the prostate cancer management options, currently there is no consensus on the standard procedure used to determine the tumor volume percentage in cases with discontinuous tumor foci. Given these controversies, we carried out a detailed characterization of molecular marker expression in cores with discontinuous foci. Our results indicated that some cores with discontinuous foci may originate from the same tumor, whereas a subset do appear to contain tumor foci from distinct clones, in keeping with results of prior studies.(26, 27) Thus, our study suggests that distinguishing the clonal origin may be helpful in assessing whether discontinuous tumor foci represent one large tumor or multiple small (and possibly clinically insignificant) tumors. As such, molecular analysis could aid in the determination of the multiclonal nature of discontinuous tumor foci, resolving dilemmas associated with cores carrying discontinuous tumor foci.

In our further analysis studying the association between molecular heterogeneity and race, we observed that the incidence of molecular heterogeneity was significantly higher in African Americans compared to Caucasian Americans. Notably, African American patients are known to present with more aggressive prostate cancer compared to the Caucasian Americans.(32) Given our findings, it would be interesting to study the clinical correlations of molecular heterogeneity in African American patients. In additional studies between the marker expression and the cancer status of biopsy cores, we observed that the expression of all four molecular markers is prostate cancer specific and mutually exclusive. Importantly, morphologically questionable cores such as atypical cores also displayed higher incidence of positive molecular markers, suggesting that molecular analysis may help eliminate ambiguities associated with prostate cancer detection on biopsies in addition to AMACR staining. Additionally, ERG was observed more in low grade cancer (Grade Group 1 and 2), whereas *ETV4* was associated with high grade cancer (Grade Group 3 and above).Further analysis on the location of tumors in the prostate gland, indicated disparity with respect to the presence of prostate cancer and molecular markers. However, it should be noted that our results could be biased due to the use of needle biopsy cores obtained from only select locations of the prostate. Further studies including the whole prostate are necessary to conclusively determine any relationship between the prostate location and the incidence of prostate cancer and molecular aberrations. A study is currently ongoing where the expression of ERG, SPINK1, *ETV1, ETV4, ETV5* and PTEN is explored in a large cohort of whole-mount radical prostatectomy samples.

As a further step, we also investigated the association of marker expression with race, and other clinical factors. We observed higher expression of SPINK1 and *ETV4* in African Americans. Additionally, SPINK1 expression and ERG+/SPINK1+ expression were found to be associated with young patient age. Of note, prior studies have implicated SPINK1 and *ETV1* in aggressive disease.(4, 7) Therefore it is interesting that we observed SPINK1 and *ETV1* to be associated with later treatment. On the other hand, *ETV1* also associated with single biopsies. However no association was identified between *ETV1* and high Gleason Grade tumors, supporting the indications that patients with *ETV1* positive tumors may have a higher chance of being directed to radical prostatectomy instead of active surveillance. Therefore additional studies including larger patient cohorts are required to establish and understand any association between the incidence of molecular markers on needle biopsies and disease outcomes. In conclusion, our study sheds light on the molecular heterogeneity and the extent of the multi clonal nature of prostate cancer at the biopsy level. Consequently, our study enables a thorough understanding of molecular heterogeneity and clonal progression of prostate cancer, facilitating the future efforts of exploring the feasibility of using molecular analysis at biopsy level as a routine step in prostate cancer management.

