## Supplementary material for "Clonal evaluation of prostate cancer molecular heterogeneity in biopsy samples by dual immunohistochemistry and dual RNA *in situ* hybridization": Tables 1-13

**Table 1: Marker status with respect to Gleason Grade group**

| **Cancer status** | **ERG+** | **SPINK1+** | **ETV1+** | **ETV4+** | **ERG+/ SPINK1+** | **ERG+/ ETV4+** | **ERG+/ETV1+** | **SPINK1+/ ETV4+** | **Negative** | ***Total number of cores*** | ***Percentage of cores showing positive expression for at least one molecular marker*** |
|---|---|---|---|---|---|---|---|---|---|---|---|
| Benign | | | | | | | | | 13 | ***13*** | ***0%*** |
| HGPIN | 2 | 5 | 2 | 3 | | | | | 111 | ***123*** | ***10%*** |
| Atypical | 5 | 5 | 1 | 1 | 1 | | | | 19 | ***32*** | ***41%*** |
| ASAP | 2 | 1 | | | | | | | 6 | ***9*** | ***33%*** |
| GG group 1 | 48 | 23 | 12 | 1 | 1 | 1 | 3 | 1 | 65 | ***155*** | ***58%*** |
| GG group 2 | 56 | 33 | 9 | | 1 | 3 | 1 | 2 | 49 | ***154*** | ***68%*** |
| GG group 3 | 8 | 9 | 1 | 2 | | | | | 24 | ***44*** | ***45%*** |
| GG group 4 | 3 | 3 | 1 | 4 | | | | | 15 | ***26*** | ***42%*** |
| GG group 5 | 9 | 3 | 2 | 2 | | | | | 29 | ***45*** | ***36%*** |
| ***Total*** | ***133*** | ***82*** | ***28*** | ***13*** | ***3*** | ***4*** | ***4*** | ***3*** | ***331*** | ***601*** | |
| ***%*** | ***22%*** | ***14%*** | ***5%*** | ***2.2%*** | ***0.5%*** | ***0.7%*** | ***0.7%*** | ***0.5%*** | ***55%*** | ***100%*** | |

**Table 2: Assessment of ERG, SPINK1, ETV1 and ETV4 in biopsy cores**

| Status of molecular marker expression | Marker status | Number of patients: Caucasian American (CA, n=67) | African American (AA, n=47) | Other (n=6) |
|---|---|---|---|---|
| Similar molecular marker expression observed with respect to ERG, SPINK1, ETV1 and ETV4 (n=76, 63%) | Tested biopsy cores are negative for ERG, SPINK1, ETV1 and ETV4 | 25 | 15 | 2 |
| | All tested biopsy cores are ERG+ | 19 | 4 | 2 |
| | All tested biopsy cores are SPINK1+ | 2 | 5 | 0 |
| | All tested biopsy cores are ETV1+ | 2 | | 0 |
| Dissimilar molecular marker expression (n=44, 37%) | ERG+ and negative cores | 3 | 2 | |
| | SPINK1+ and negative cores | 3 | 7 | 1 |
| | ETV1+ and negative cores | 2 | | |
| | ETV4+ and negative cores | 1 | | |
| | ERG+ and SPINK1+ cores | 2 | 5 | |
| | SPINK1+ and ETV1+ cores | | 1 | |
| | SPINK1+ and ETV4+ cores | | 1 | |
| | ERG+ and ETV4+ cores | 1 | 3 | |
| | ERG+ and ETV1+ cores | 7 | 1 | 1 |
| | ETV1+ and ETV4+ cores | | 1 | |
| | ERG+, SPINK1+ and ETV4+ cores | | 1 | |
| | ERG+, SPINK1+ and ETV1+ cores | | 1 | |
| | Total number of patients | 67 | 47 | 6 |

"ERG+ and negative cores" refer to cases where ERG+ biopsy cores were observed along with biopsy cores negative for ERG, SPINK1, ETV1 and ETV5. Similarly, "SPINK1+ and negative cores", "ETV1+ and negative cores" and "ETV4+ and negative cores" refer to cases where biopsy cores positive for the corresponding molecular marker were observed along with biopsy cores negative for ERG, SPINK1, ETV1 and ETV5. "ERG+ and SPINK1+ cores", "SPINK1+ and ETV1+ cores", "SPINK1+ and ETV4 cores", "ERG+ and ETV4+ cores", "ERG+ and ETV1+ cores", "ETV1+ and ETV4+ cores","ERG+, SPINK1+ and ETV4+ cores","ERG+, SPINK1+ and ETV1+ cores" refer to cases with biospy cores showing multiple molecualr marker positivity. For example, "ERG+ and SPINK1+ cores" refers to cases where both ERG+ and SPINK1+ biospy cores were observed.

**Table S1: The pateint cohort**

| Case number | Patient Status as of May-2019 | Race | Patient Age at the time of study biopsy | Family history of prostate cancer | Initial PSA (ng/mL) | Time between study biopsy and initial PSA | number of biopsies before study biopsy | number of biopsies after study biopsy | total number of biopsies including the study biopsy | Subsequent treatment used | Time from the study biopsy to the subsequent treatment | Last PSA (ng/mL) as of May 2019 | Time from treatement to the last PSA |
|---|---|---|---|---|---|---|---|---|---|---|---|---|---|
| 1 | Alive | African American | 59 | positive | 8.4 | <1 month | | | 1 | n/a | Not available | 9.7 | Not available |
| 2 | Alive | African American | 72 | negative | 11.5 | <1 month after study biopsy | | | 1 | RT | 3 | 0.4 | 26 |
| 3 | Alive | African American | 75 | negative | 5.8 | 2 | 1 | | 2 | RP | 1 | <0.1 | 15 |
| 4 | Alive | African American | 65 | negative | 8.8 | 1 | | | 1 | n/a | Not available | 13.5 | Not available |
| 5 | Alive | Caucasian American | 58 | positive | 6.5 | 1 | | | 1 | n/a | Not available | 4.6 | Not available |
| 6 | Alive | Caucasian American | 81 | negative | 35.3 | <1 month | | | 1 | HT | <1 month | 4.7 | 34 |
| 7 | Alive | Caucasian American | 66 | negative | 11.4 | 2 | 3 | | 4 | n/a | Not available | 1.9 | Not available |
| 8 | Alive | Caucasian American | 80 | negative | 7.2 | <1 month | 1 | | 2 | HT | 4 | <0.1 | 27 |
| 9 | Alive | African American | 71 | negative | 27.6 | <1 month | | | 1 | RP | 4 | 0.5 | 5 |
| 10 | Alive | African American | 73 | positive | 4.9 | 4 | 2 | | 3 | n/a | Not available | 4.2 | Not available |
| 11 | Alive | African American | 57 | positive | 13.4 | <1 month after study biopsy | | | 1 | RP/RT | 7 | <0.1 | 13 |
| 12 | Alive | African American | 53 | negative | 4.5 | <1 month | | | 1 | RP | 2 | <0.1 | 30 |
| 13 | Alive | African American | 55 | positive | 11.6 | 1 | | | 1 | n/a | Not available | o follow up PS. | Not available |
| 14 | Alive | Caucasian American | 62 | negative | 3.8 | 2 | | | 1 | n/a | Not available | o follow up PS. | Not available |
| 15 | Alive | other | 56 | positive | 6 | 6 | | | 1 | RP | 4 | 0.3 | 25 |
| 16 | Alive | Caucasian American | 73 | negative | 0.5 | 15 | | | 1 | n/a | Not available | o follow up PS. | Not available |
| 17 | Alive | Caucasian American | 59 | negative | 0.026 | <1 month after study biopsy | | | 1 | n/a | Not available | 1.93 | Not available |
| 18 | Alive | African American | 73 | negative | 6.3 | <1 month | | 1 | 2 | RT/HT | 12 | 0.2 | 20 |
| 19 | Alive | African American | 58 | negative | 18.5 | 1 | 2 | | 3 | RP | 1 | 0.3 | 29 |
| 20 | Alive | African American | 63 | negative | 5.2 | 2 | | | 1 | RT | Not available | 7.9 | Not available |
| 21 | Alive | African American | 76 | negative | 12.9 | 1 | | | 1 | n/a | Not available | <0.1 | Not available |
| 22 | Alive | other | 71 | negative | 5.9 | 1 | | 1 | 2 | RP | 14 | <0.1 | 15 |
| 23 | Alive | African American | 68 | negative | 6.4 | 1 | | | 1 | RP | 4 | <0.1 | 17 |
| 24 | Alive | African American | 55 | negative | 6.5 | 1 | | | 1 | RP/RT/HT | 1 | <0.1 | 28 |
| 25 | Alive | Caucasian American | 62 | negative | 5.7 | 1 | | | 1 | RP | 3 | <0.1 | 26 |
| 26 | Alive | other | 78 | negative | 102.2 | <1 month after study biopsy | | | 1 | RP/HT/RT | 11 | <0.1 | 19 |
| 27 | Alive | African American | 74 | negative | 4.4 | 1 | | | 1 | RP | 1 | <0.1 | 28 |
| 28 | Alive | Caucasian American | 64 | negative | 5 | <1 month | | | 1 | n/a | Not available | 3.8 | Not available |
| 29 | Alive | Caucasian American | 65 | negative | 20.2 | <1 month | | | 1 | n/a | Not available | <0.1 | Not available |
| 30 | Alive | African American | 48 | positive | 1.9 | <1 month | 1 | 1 | 3 | n/a | Not available | 6.35 | Not available |
| 31 | Alive | Caucasian American | 74 | negative | 5.4 | 3 | | | 1 | RP | <1 month | <0.1 | 32 |
| 32 | Alive | African American | 65 | negative | 29.2 | <1 month | 1 | | 2 | RT/HT | <1 month | <0.1 | 31 |
| 33 | Alive | Caucasian American | 80 | negative | 4.9 | <1 month | | | 1 | n/a | Not available | 0.6 | Not available |
| 34 | Alive | African American | 70 | negative | 5.2 | <1 month | | | 1 | RP | 2 | <0.1 | 18 |
| 35 | Alive | Caucasian American | 80 | negative | 5.2 | 2 | | 1 | 2 | RT/HT | 16 | <0.1 | 13 |
| 36 | Alive | African American | 60 | negative | 4 | 1 | | | 1 | RP | 6 | <0.1 | 13 |
| 37 | Alive | African American | 45 | negative | 5.4 | 1 | | | 1 | RT | 11 | 1.4 | 21 |
| 38 | Alive | African American | 53 | negative | 16.4 | <1 month | | | 1 | HT | <1 month | 16.3 | 33 |
| 39 | Alive | African American | 50 | negative | 6.8 | 1 | | | 1 | RP | 2 | <0.1 | 24 |
| 40 | Alive | African American | 64 | negative | 1.7 | <1 month | | | 1 | RP | 2 | <0.1 | 28 |
| 41 | Alive | Caucasian American | 74 | negative | 4.1 | 1 | | | 1 | RP | 3 | <0.1 | 27 |
| 42 | Alive | African American | 66 | negative | 9.7 | <1 month after study biopsy | 1 | | 2 | RT | 2 | 1.2 | 22 |
| 43 | Alive | African American | 65 | negative | 8.2 | <1 month | | | 1 | n/a | Not available | 0.3 | Not available |
| 44 | Alive | Caucasian American | 71 | negative | 2.8 | <1 month | | 1 | 2 | RP | 14 | 8.4 | 14 |
| 45 | Alive | Caucasian American | 67 | negative | 13.4 | <1 month | | | 1 | n/a | Not available | 12.2 | Not available |
| 46 | Alive | Caucasian American | 86 | negative | 3.6 | 2 | | | 1 | n/a | Not available | 0.3 | Not available |
| 47 | Alive | Caucasian American | 62 | positive | 5.2 | 1 | | | 1 | RP | 4 | <0.1 | 18 |
| 48 | Alive | Caucasian American | 64 | negative | 5.1 | 2 | 1 | | 2 | n/a | Not available | 5.2 | Not available |
| 49 | Alive | Caucasian American | 74 | negative | 5.1 | <1 month | | | 1 | RT | 3 | 0.9 | 28 |
| 50 | Alive | African American | 60 | negative | 3.8 | 6 | | | 1 | RP | 1 | 3.8 | 25 |
| 51 | Alive | African American | 63 | negative | 7.3 | 1 | | | 1 | RT | 27 | 0.7 | 3 |
| 52 | Alive | Caucasian American | 84 | positive | 21.5 | 2 | | | 1 | RT/HT | <1 month | 27.2 | 6 |
| 53 | Alive | Caucasian American | 60 | negative | 6 | <1 month | | | 1 | RP | 1 | <0.1 | 30 |
| 54 | Alive | other | 55 | positive | 16.4 | <1 month | | | 1 | RP | 8 | 16.4 | 1 |
| 55 | Alive | Caucasian American | 71 | negative | 1 | 1 | 2 | | 3 | n/a | Not available | o follow up PS. | Not available |
| 56 | Alive | African American | 61 | negative | 9.8 | 1 | | | 1 | RT | Not available | <0.1 | Not available |
| 57 | Alive | Caucasian American | 63 | positive | 4.2 | 2 | | 1 | 2 | n/a | Not available | 5.9 | Not available |
| 58 | Alive | African American | 65 | negative | 4.8 | <1 month | | | 1 | RP | 8 | <0.1 | 19 |
| 59 | Alive | Caucasian American | 68 | negative | 5.4 | <1 month | | | 1 | n/a | Not available | o follow up PS. | Not available |
| 60 | Alive | African American | 53 | negative | 4.2 | 2 | 1 | | 2 | RP | 2 | <0.1 | 30 |
| 61 | Alive | Caucasian American | 50 | negative | 13.4 | 1 | | | 1 | RP | 2 | <0.1 | 29 |
| 62 | Alive | Caucasian American | 69 | negative | 12.9 | <1 month | 7 | | 8 | n/a | Not available | 16.0 | Not available |
| 63 | Alive | Caucasian American | 54 | negative | 7.8 | <1 month | | | 1 | RP | <1 month | <0.1 | 29 |
| 64 | Alive | Caucasian American | 61 | positive | 1810.8 | <1 month | | | 1 | HT | <1 month | 0.3 | 30 |
| 65 | Alive | African American | 70 | negative | 11.1 | 1 | 1 | | 2 | RP | 1 | <0.1 | 18 |
| 66 | Alive | Caucasian American | 69 | negative | 4.6 | <1 month | | | 1 | RP | 2 | <0.1 | 29 |
| 67 | Alive | African American | 70 | negative | 10.4 | 2 | | | 1 | n/a | Not available | 0.6 | Not available |
| 68 | Alive | African American | 59 | negative | 4.5 | <1 month | | | 1 | RP | 7 | <0.1 | 15 |
| 69 | Alive | Caucasian American | 65 | negative | 7.5 | <1 month | 2 | | 3 | RT | 2 | 0.7 | 26 |
| 70 | Alive | African American | 75 | negative | 7.1 | 3 | 1 | | 2 | RT/HT | 16 | <0.1 | 11 |
| 71 | Alive | Caucasian American | 68 | negative | 5.3 | 3 | | 1 | 2 | n/a | Not available | 3.0 | Not available |
| 72 | Deceased | Caucasian American | 80 | negative | 12.4 | <1 month | 3 | | 4 | n/a | Not available | 18.1 | Not available |
| 73 | Alive | Caucasian American | 66 | positive | 0.8 | 8 | 3 | 1 | 5 | n/a | Not available | 0.9 | Not available |
| 74 | Alive | African American | 74 | negative | 5.9 | 5 | | 2 | 3 | n/a | Not available | <0.1 | Not available |
| 75 | Alive | African American | 80 | negative | 18.6 | <1 month | | | 1 | n/a | Not available | 13.9 | Not available |
| 76 | Alive | African American | 65 | positive | 4.5 | <1 month | | | 1 | RT | 7 | <0.1 | 23 |
| 77 | Alive | Caucasian American | 74 | ormation not availal | 9.1 | <1 month | 2 | | 3 | n/a | Not available | 12.4 | Not available |
| 78 | Alive | Caucasian American | 66 | negative | 5.5 | 14 | 1 | | 2 | RP | 3 | <0.1 | 23 |
| 79 | Alive | African American | 68 | negative | 5.5 | 4 | 1 | 1 | 3 | n/a | Not available | 7.4 | Not available |
| 80 | Alive | Caucasian American | 65 | positive | 4.9 | <1 month | | | 1 | RP | 3 | 4.9 | 22 |
| 81 | Alive | African American | 64 | negative | 1.1 | <1 month | 1 | | 2 | n/a | Not available | 1.1 | Not available |
| 82 | Alive | Caucasian American | 72 | negative | 8.7 | 6 | 2 | | 3 | n/a | Not available | 10.5 | Not available |
| 83 | Alive | Caucasian American | 81 | negative | 9.4 | <1 month after study biopsy | | | 1 | RP | 1 | 0.3 | 27 |
| 84 | Alive | African American | 62 | negative | 4 | <1 month | | | 1 | RT | 7 | o follow up PS. | Not available |
| 85 | Alive | African American | 50 | positive | 6.1 | 7 | | | 1 | RP | 2 | <0.1 | 1 |
| 86 | Alive | African American | 62 | positive | 4.2 | 5 | | | 1 | n/a | Not available | 5.4 | Not available |
| 87 | Alive | African American | 72 | negative | 5.7 | <1 month | | 1 | 2 | n/a | Not available | 7.7 | Not available |
| 88 | Alive | Caucasian American | 73 | negative | 11.4 | 1 | 2 | | 3 | RT | <1 month | <0.1 | 30 |
| 89 | Alive | Caucasian American | 71 | negative | 6.8 | <1 month | | | 1 | n/a | Not available | o follow up PS. | Not available |
| 90 | Alive | Caucasian American | 51 | positive | 4.2 | 1 | | 2 | 3 | RP | 9 | <0.1 | 22 |
| 91 | Alive | other | 75 | negative | 5.3 | 2 | | | 1 | n/a | Not available | o follow up PS. | Not available |
| 92 | Alive | Caucasian American | 73 | negative | 3.3 | 2 | | | 1 | n/a | Not available | 2.8 | Not available |
| 93 | Alive | Caucasian American | 58 | positive | 12.1 | 1 | | | 1 | RP/RT | 1 | <0.1 | 33 |
| 94 | Alive | Caucasian American | 64 | negative | 4.8 | 3 | | | 1 | RT | Not available | 0.6 | Not available |
| 95 | Alive | Caucasian American | 61 | positive | 5.4 | 1 | | | 1 | RP | 3 | <0.1 | 27 |
| 96 | Alive | Caucasian American | 81 | negative | 1.3 | 5 | | 1 | 2 | n/a | Not available | 0.4 | Not available |
| 97 | Alive | Caucasian American | 53 | negative | 3.1 | 2 | | | 1 | RP | 3 | <0.1 | 28 |
| 98 | Alive | Caucasian American | 54 | negative | 0.7 | 1 | 2 | 1 | 4 | n/a | Not available | 0.9 | Not available |
| 99 | Alive | other | 75 | negative | 4.5 | 2 | | | 1 | n/a | Not available | 5.9 | Not available |
| 100 | Alive | Caucasian American | 77 | negative | 5.6 | 3 | | 1 | 2 | n/a | Not available | 18.3 | Not available |
| 101 | Alive | Caucasian American | 61 | positive | 4.7 | 2 | | | 1 | RP | 2 | <0.1 | 28 |
| 102 | Alive | Caucasian American | 54 | negative | 0.8 | <1 month | 3 | | 4 | n/a | Not available | 0.9 | Not available |
| 103 | Alive | Caucasian American | 44 | negative | 4.5 | 3 | | | 1 | RP | 2 | <0.1 | 28 |
| 104 | Alive | Caucasian American | 81 | negative | 15.1 | 2 | | | 1 | RP | 2 | 1.5 | 29 |
| 105 | Alive | Caucasian American | 71 | negative | 5.2 | 1 | | | 1 | n/a | Not available | 4.0 | Not available |
| 106 | Alive | Caucasian American | 53 | negative | 11.2 | 1 | | | 1 | RP | 1 | <0.1 | 1 |
| 107 | Alive | Caucasian American | 76 | negative | 11.8 | 3 | | | 1 | n/a | Not available | 11.0 | Not available |
| 108 | Alive | Caucasian American | 70 | negative | 3.7 | <1 month | | | 1 | n/a | Not available | <0.1 | Not available |
| 109 | Alive | Caucasian American | 69 | negative | 4.9 | 1 | | | 1 | RP/RT | 2 | 0.3 | 29 |
| 110 | Alive | Caucasian American | 73 | negative | 3.2 | 1 | | | 1 | RP | 1 | <0.1 | 30 |
| 111 | Alive | Caucasian American | 62 | negative | 5.1 | 6 | | 1 | 2 | n/a | Not available | 2.0 | Not available |
| 112 | Alive | African American | 58 | positive | 2.8 | 3 | | | 1 | RP | 1 | <0.1 | 29 |
| 113 | Alive | Caucasian American | 51 | negative | 4.3 | 2 | | | 1 | RP | 3 | <0.1 | 24 |
| 114 | Alive | Caucasian American | 65 | negative | 4.2 | <1 month | | | 1 | RP | 2 | <0.1 | 28 |
| 115 | Alive | Caucasian American | 63 | negative | 8 | 3 | | | 1 | RP | 3 | <0.1 | 29 |
| 116 | Alive | African American | 67 | negative | 4.4 | 4 | | | 1 | RP | 3 | <0.1 | 28 |
| 117 | Alive | Caucasian American | 55 | negative | 4.9 | 1 | | 1 | 2 | RP | 15 | <0.1 | 15 |
| 118 | Alive | Caucasian American | 81 | positive | 12.5 | <1 month | | | 1 | n/a | Not available | 6.1 | Not available |
| 119 | Alive | Caucasian American | 71 | negative | 4.2 | 1 | | | 1 | n/a | Not available | 3.0 | Not available |
| 120 | Alive | African American | 67 | negative | 7.1 | 1 | | | 1 | RP | 3 | <0.1 | 28 |

**Table S2: The number of cores collected from each location of the prostate**

| Prostate location from where the needle core tissue was extracted | Number of cores | Percentage | |
|---|---|---|---|
| RLB | 39 | 6% | Needle biopsy cores collected from standard 12 core biopsy locations (n=572) |
| RLM | 50 | 8% | |
| RLA | 48 | 8% | |
| RB | 41 | 7% | |
| RM | 49 | 8% | |
| RA | 54 | 9% | |
| LB | 39 | 6% | |
| LM | 46 | 8% | |
| LA | 54 | 9% | |
| LLB | 45 | 7% | |
| LLM | 47 | 8% | |
| LLA | 60 | 10% | |
| Right medial anterior | 1 | 0% | Needle biopsy cores collected from other prostate locations (n=29) |
| Right medial anterior perineal | 1 | 0% | |
| Right medial posterior | 1 | 0% | |
| Right lateral anterior | 2 | 0% | |
| Right anterior | 9 | 1% | |
| Right lateral posterior perineal | 1 | 0% | |
| Rightmost lateral anterior perineal | 1 | 0% | |
| Rightmost lateral posterior perineal | 1 | 0% | |
| Left anterior | 9 | 1% | |
| Left medial posterior | 1 | 0% | |
| Left lateral posterior | 1 | 0% | |
| Leftmost lateral anterior | 1 | 0% | |
| ***Total number of cores*** | ***601*** | ***100%*** | |

**Table S3: The cancer status of the needle biopsy cores obtained from different locations of the prostate**

| Tumor Location in prostate | Cancer status | | | | | | | | | Total number of cores | Percentage of cores showing GG 1-5 cancer, Atypical lesion or ASAP |
|---|---|---|---|---|---|---|---|---|---|---|---|
| | **GG 1** | **GG 2** | **GG 3** | **GG 4** | **GG 5** | **benign** | **HGPIN** | **ASAP** | **Atypical** | | |
| RLB | 10 | 10 | 1 | 1 | 2 | | 11 | | 4 | ***39*** | **72%** |
| RLM | 15 | 12 | 3 | 1 | 3 | 1 | 11 | 1 | 3 | ***50*** | **76%** |
| RLA | 17 | 9 | 5 | 1 | 5 | 1 | 7 | 2 | 1 | ***48*** | **83%** |
| RB | 7 | 11 | 4 | 1 | 3 | | 13 | 1 | 1 | ***41*** | **68%** |
| RM | 12 | 10 | 3 | 2 | 4 | 2 | 14 | | 2 | ***49*** | **67%** |
| RA | 18 | 13 | 2 | 2 | 3 | 4 | 9 | | 3 | ***54*** | **76%** |
| LB | 6 | 11 | 5 | 2 | 4 | 1 | 8 | | 2 | ***39*** | **77%** |
| LM | 10 | 14 | 3 | 4 | 4 | 2 | 7 | 2 | | ***46*** | **80%** |
| LA | 15 | 17 | 5 | 3 | 5 | 1 | 5 | 1 | 2 | ***54*** | **89%** |
| LLB | 10 | 10 | 4 | 4 | 2 | 1 | 9 | | 5 | ***45*** | **78%** |
| LLM | 11 | 14 | 5 | 2 | 4 | | 8 | | 3 | ***47*** | **83%** |
| LLA | 19 | 14 | 3 | 2 | 4 | | 12 | 2 | 4 | ***60*** | **80%** |
| ***Total*** | ***150*** | ***145*** | ***43*** | ***25*** | ***43*** | ***13*** | ***114*** | ***9*** | ***30*** | ***572*** | |
| ***%*** | ***26%*** | ***25%*** | ***8%*** | ***4%*** | ***8%*** | ***2%*** | ***20%*** | ***2%*** | ***5%*** | ***100%*** | |

**Table S4: The expression of molecular markers in cores obtained from different locations of the prostate**

| **Prostate core location** | **ERG+** | **SPINK1+** | **ETV1+** | **ETV4+** | **ERG+/ SPINK1+** | **ERG+/E TV4+** | **ERG+/E TV1+** | **SPINK1+/ ETV4+** | **Molecular Marker Positive Cores** | **Molecular Marker Negative Cores** | ***Percenatge of cores showing positive expression for at least one molecular marker*** |
|---|---|---|---|---|---|---|---|---|---|---|---|
| RLB | 7 | 8 | 3 | 1 | | 1 | | | 20 | 19 | **51%** |
| RLM | 7 | 9 | 5 | 1 | | | 1 | | 23 | 27 | **46%** |
| RLA | 8 | 9 | 3 | 1 | 1 | 1 | 1 | | 24 | 24 | **50%** |
| RB | 8 | 5 | 2 | 1 | | | | | 16 | 25 | **39%** |
| RM | 9 | 4 | 2 | 2 | 1 | | | | 18 | 31 | **37%** |
| RA | 12 | 6 | 5 | | | | | | 23 | 31 | **43%** |
| LB | 5 | 5 | 1 | 2 | | | | | 13 | 26 | **33%** |
| LM | 14 | 7 | 1 | 1 | 1 | 1 | | | 25 | 21 | **54%** |
| LA | 16 | 7 | 1 | 1 | | | | | 25 | 29 | **46%** |
| LLB | 12 | 8 | 1 | 1 | | | 1 | 1 | 24 | 21 | **53%** |
| LLM | 13 | 7 | 1 | 1 | | 1 | 1 | 1 | 25 | 22 | **53%** |
| LLA | 19 | 5 | | 1 | | | | | 25 | 35 | **42%** |
| ***Total*** | ***130*** | ***80*** | ***25*** | ***13*** | ***3*** | ***4*** | ***4*** | ***2*** | ***261*** | ***311*** | **46%** |
| **%** | **23%** | **14%** | **4%** | **2%** | **1%** | **1%** | **1%** | **0%** | **46%** | **54%** | |

| Table S5: The incidence of ERG, SPINK1, ETV1 and ETV4 in the patient cohort | | | | |
|---|---|---|---|---|
| **Marker status** | **Marker expression** | **Caucasian American (CA, n=67)** | **African American (AA, n=47)** | **Other (n=6)** |
| | ERG+ | 22 | 6 | 2 |
| Expression of a single marker (n=53) | SPINK1+ | 5 | 12 | 1 |
| | ETV1+ | 4 | | 0 |
| | ETV4+ | 1 | | |
| | ERG+/SPINK1+ | 2 | 5 | |
| | SPINK1+/ ETV1+ | | 1 | |
| Expression of two different markers (n=23) | SPINK1+/ETV4+ | | 1 | |
| | ERG+/ETV4+ | 1 | 3 | |
| | ERG+/ETV1+ | 7 | 1 | 1 |
| | ETV1+/ETV4+ | | 1 | |
| Expression of three different markers (n=2) | ERG+, SPINK1+, ETV4+ | | 1 | |
| | ERG+, SPINK1+, ETV1+ | | 1 | |
| Negative for all tested markers | | 25 | 15 | 2 |

**Table S6: The association of molecular marker expression with the patient age**

| Molecular Marker | Mean age of the cases negative for all the markers (SD) | Mean age of the cases positive for one or more markers(SD) | p-value |
|---|---|---|---|
| ERG | 67.30 (9.18) | 64.08 (8.76) | 0.06 |
| SPINK1 | 67.08 (8.64) | 62.18 (9.74) | **0.01** |
| ETV1 | 66.26 (8.87) | 63.81 (10.59) | 0.32 |
| ETV4 | 65.81 (9.23) | 67.62 (7.39) | 0.59 |
| ERG/ETV4 | 65.89 (9.18) | 67.25 (7.50) | 0.77 |
| ERG/ETV1 | 66.33 (9.02) | 60.38 (9.02) | 0.07 |
| ERG/SPINK1 | 66.39 (8.90) | 57.33 (9.54) | **0.02** |

**Table S7: The association of molecular marker expression with the number of biopsies**

| Molecular Marker | Biopsy group | No of cases with the marker negative | No of cases with the marker positive | p-value |
|---|---|---|---|---|
| ERG | Multiple biopsies(>1) | 26 (37.7%) | 13 (25.5%) | |
| | One Biopsy | 43 (62.3%) | 38 (74.5%) | 0.23 |
| SPINK1 | Multiple biopsies(>1) | 29 (31.5%) | 10 (35.7%) | |
| | One Biopsy | 63 (68.5%) | 18 (64.3%) | 0.85 |
| ETV1 | Multiple biopsies(>1) | 39 (37.5%) | 0 (0.0%) | |
| | One Biopsy | 65 (62.5%) | 8 (100.0%) | **0.01** |
| ETV4 | Multiple biopsies(>1) | 36 (32.1%) | 3 (37.5%) | |
| | One Biopsy | 76 (67.9%) | 5 (62.5%) | 1 |
| ERG/ETV4 | Multiple biopsies(>1) | 38 (32.8%) | 1 (25.0%) | |
| | One Biopsy | 78 (67.2%) | 3 (75.0%) | 1 |
| ERG/ETV1 | Multiple biopsies(>1) | 39 (34.8%) | 0 (0.0%) | |
| | One Biopsy | 73 (65.2%) | 8 (100.0% | 0.10 |
| ERG/SPINK1 | Multiple biopsies(>1) | 38 (33.3%) | 1 (16.7%) | |
| | One Biopsy | 76 (66.7%) | 5 (83.3%) | 0.69 |

**Table S8: The association of molecular marker expression with intial PSA value**

| Molecular Marker | mean initial PSA value in cases where marker was negative (SD) | mean initial PSA value in cases where marker was positive (SD) | p-value |
|---|---|---|---|
| ERG | 9.19 (13.18) | 42.22 (252.63) | 0.28 |
| SPINK1 | 26.43 (188.14) | 12.69 (19.13) | 0.70 |
| ETV1 | 25.56 (177.08) | 8.05 (7.20) | 0.69 |
| ETV4 | 24.10 (170.67) | 10.93 (7.26) | 0.83 |
| ERG/ETV4 | 23.74 (167.70) | 8.43 (2.35) | 0.86 |
| ERG/ETV1 | 24.46 (170.65) | 5.94 (3.05) | 0.76 |
| ERG/SPINK1 | 24.09 (169.15) | 6.73 (2.14) | 0.80 |

**Table S9: The association of molecular marker expression with last PSA value**

| Molecular Marker | mean of the last PSA value in cases where marker was negative (SD) | mean of the last PSA value in cases where marker was positive (SD) | p-value |
|---|---|---|---|
| ERG | 5.74 (6.46) | 4.30 (4.62) | 0.33 |
| SPINK1 | 5.03 (5.77) | 5.86 (6.28) | 0.65 |
| ETV1 | 4.95 (5.20) | 8.25 (11.59) | 0.23 |
| ETV4 | 5.06 (5.69) | 7.22 (8.57) | 0.48 |
| ERG/ETV4 | 5.20 (5.87) | 5.05 (6.58) | 0.97 |
| ERG/SPINK1 | 5.33 (5.88) | 1.00 (0.57) | 0.31 |

**Table S10: The association between the molecular marker expression and the presence of subsequent treatment**

| Molecualr Marker | p-value | HR | CI |
|---|---|---|---|
| ERG+ | 0.67 | 1.1069 | 0.6921 1.77 |
| SPINK1+ | **0.01** | 1.9777 | 1.191 3.285 |
| ETV1+ | **0.00** | 2.5074 | 1.329 4.731 |
| ETV4+ | 0.46 | 1.372 | 0.5917 3.181 |
| ERG+ with other markers negative | **0.05** | 0.5504 | 0.3064 0.9885 |
| SPINK1+ with other markers negative | 0.05 | 1.7829 | 0.9916 3.206 |
| ETV1+ with other markers negative | 0.92 | 1.0775 | 0.2636 4.404 |
| ERG+/ETV1+ | **0.00** | 3.373 | 1.561 7.287 |
| ERG+/ETV4+ | 0.31 | 1.8359 | 0.5716 5.896 |
| ERG+/SPINK1+ | 0.31 | 1.6076 | 0.646 4 |
| All markers negative | 0.09 | 0.6447 | 0.3862 1.076 |

**Table S11: The association of molecular marker expression with subsequent radical prostatectomy**

| Molecualr Marker | p-value | HR | CI |
|---|---|---|---|
| ERG+ | 0.27 | 1.3678 | 0.7879 2.374 |
| SPINK1+ | 0.15 | 1.5823 | 0.8537 2.933 |
| ETV1+ | **<0.01** | 3.2012 | 1.621 6.323 |
| ETV4+ | 0.38 | 1.5185 | 0.6003 3.841 |
| ERG+ with other markers negative | 0.10 | 0.5555 | 0.2781 1.109 |
| SPINK1+ with other markers negative | 0.77 | 1.125 | 0.5062 2.5 |
| ETV1+ with other markers negative | 0.77 | 0.7393 | 0.102 5.357 |
| ERG+/ETV1+ | **<0.01** | 4.661 | 2.103 10.33 |
| ERG+/ETV4+ | 0.59 | 1.482 | 0.3577 6.14 |
| ERG+/SPINK1+ | 0.31 | 1.6993 | 0.6117 4.72 |
| All markers negative | 0.19 | 0.6656 | 0.3634 1.219 |

**Table S12: The association of molecular marker expression with subsequent radiation**

| Molecular Marker | p-value | HR | CI |
|---|---|---|---|
| ERG+ | 0.807 | 0.6086 | 0.352<br>3.825 |
| SPINK1+ | 0.0835 | 3.0727 | 0.8619<br>10.95 |
| ETV1+ | 0.998 | 3.504E-08 | 0<br>Inf |
| ETV4+ | 0.636 | 1.6476 | 0.208<br>13.05 |
| ERG+ with other markers negative | 0.683 | 0.7578 | 0.1999<br>2.873 |
| SPINK1+ with other markers negative | 0.0999 | 3.0895 | 0.8058<br>11.84 |
| ETV1+ with other markers negative | 0.999 | 3.931E-08 | 0<br>Inf |
| ERG+/ETV1+ | 0.999 | 3.673E-08 | 0<br>Inf |
| ERG+/ETV4+ | 0.0806 | 7.058 | 0.7881<br>63.2 |
| ERG+/SPINK1+ | 0.367 | 2.6241 | 0.3227<br>21.34 |
| All markers negative | 0.35 | 0.5291 | 0.1391<br>2.012 |

**Table S13: The association of molecular marker expression with the duration of time occurred between the study biopsy and the subsequent treatment**

| Molecular Marker | mean value for the time (months) in cases where marker was negative (SD) | mean value for the time (months) in cases where marker was positive (SD) | p-value |
|---|---|---|---|
| ERG | 5.18 (6.26) | 3.30 (2.53) | 0.117 |
| SPINK1 | 4.09 (4.31) | 4.77 (6.21) | 0.595 |
| ETV1 | 4.75 (5.33) | 2.17 (1.09) | 0.1 |
| ETV4 | 4.38 (5.15) | 3.50 (1.97) | 0.68 |
| ERG expression when other markers are negative | 4.47 (5.38) | 3.64 (2.73) | 0.58 |
| SPINK1 expression when other markers are negative | 4.14 (4.17) | 4.96 (7.44) | 0.58 |
| ETV1 expression when other markers are negative | 4.42 (4.98) | 0.50 (0.00) | 0.274 |
| ERG/ETV1 | 4.56 (5.21) | 2.38 (0.74) | 0.244 |
| ERG/ETV4 | 4.39 (5.04) | 2.33 (1.15) | 0.485 |
| ERG/SPINK1 | 4.27 (5.03) | 4.70 (4.27) | 0.854 |
| All markers negative | 3.74 (4.47) | 5.79 (5.93) | 0.126 |
